# Convergent evolution of bird-mammal shared characteristics for adapting to nocturnality

**DOI:** 10.1101/403139

**Authors:** Yonghua Wu, Haifeng Wang

## Abstract

The diapsid lineage (birds) and synapsid lineage (mammals), share a suite of functionally similar characteristics (e.g., endothermy) that are considered to be a result of their convergent evolution, but the candidate selections leading to this convergent evolution are still under debates. Here, we used a newly developed molecular phyloecological approach to reconstruct the diel activity pattern of the common ancestors of living birds. Our results strongly suggest that they had adaptations to nocturnality during their early evolution, which is remarkably similar to that of ancestral mammals. Given their similar adaptation to nocturnality, we propose that the shared traits in birds and mammals may have evolved as a result of the convergent evolution of their early ancestors adapting to ecological factors (e.g., low ambient temperature) associated with nocturnality. Finally, a conceptually unifying ecological model on the evolution of endothermy in diverse organisms with an emphasis on low ambient temperature is proposed. We reason that endothermy may evolve as an adaptive strategy to enable organisms to effectively implement various life cycle activities under relatively low-temperature environments. In particular, a habitat shift from high-temperature to relatively low-temperature environments is identified as a common factor underlying the evolution of endothermy.

## Introduction

Extant birds and mammals share a number of highly similar characteristics, including but not limited to, enhanced hearing, vocal communication, endothermy, insulation, shivering, respiratory turbinates, high basal metabolism, grinding, sustained activity, four chambered heart, high blood pressure and intensive parental care ^1-8^. These bird-mammal shared characteristics (BMSC) are considered to have evolved convergently in the two groups ^4^, while the selection pressures that underlie their convergent evolution are still under debates. To date, many candidate selections, which are not necessarily mutually exclusive, have been proposed ^9-11^ and popular views suggest that BMSC (e.g., endothermy) evolved primarily for increased sustained activity ^1^ and enhanced parental care ^4,8^. On the other hand, the traditional view, the thermal niche expansion model, invokes nocturnality as an initial driver for the evolution of endothermy in ancestral mammals that would not necessarily featured by high resting metabolic rate typical of most living mammals ^2,7^. Indeed, early mammals are widely accepted as having been nocturnal ^12,13^, and it has been hypothesized that the evolution of endothermy may have allowed them to overcome ambient low temperature constraints to be active at night, while diurnal ectothermic reptiles essentially depend on sunlight to increase their body temperature to operational levels ^12,14^. The nocturnality hypothesis is criticized as being suitable only to mammals ^1^, partly because so far only the nocturnality of the synapsid lineages (e.g., early mammals) is well known, while the possible nocturnal activity of the diapsid lineages (e.g., birds), especially during their early evolution, is unknown.

The diel activity pattern of ancestral birds has received less attention. To date, there are only two relevant studies based on behavior or morphology analyses. A maximum likelihood reconstruction based on behavioral data suggests the diurnality of the common ancestor of living birds (CALB) ^15^. This study classifies each species as either diurnal or nocturnal without consideration of other behavioral possibilities (e.g., crepuscular or cathemeral) or behavioral complexity, for instance, the widespread existence of the occasional nocturnality of typically diurnal birds ^16^, and hence this result may be less accurate. Another study uses a phylogenetic discriminant analysis to reconstruct the diel activity patterns of four fossil basal birds (*Archaeopteryx lithographica, Confuciusornis sanctus, Sapeornis chaoyangensis, Yixianornis grabaui*) based on eye morphology data, and the result demonstrates their diurnality ^17^. The phylogenetic discriminant analysis approach is criticized for its incapability to distinguish diurnality from cathemerality (i.e., active both day and night) ^18,19^, and hence uncertainty remains concerning the diel activity patterns of these basal birds. Given the uncertainties of the results, the diel activity pattern of ancestral birds remains unclear.

The recent development of a molecular phyloecological (MPE) approach allows us to reconstruct the diel activity pattern of ancestral taxa using molecular data ^13,20-22^. The MPE approach has been applied to analyze the diel activity patterns of birds and mammals, and the results demonstrate its sensitivity to distinct animals with different diel activity patterns, showing that nocturnality, diurnality and cathemerality are featured by enhanced positive selection of dim-light vision genes (rod-expressed genes), bright-light vision genes (cone-expressed genes) and both, respectively ^13,20,22^. In this study, we employed the MPE approach to reconstruct the early evolution of the diel activity patterns in diapsid lineages. Our results reveal that diapsid lineages exhibit similar nocturnality adaption to that of synapsid lineages. Our findings provide insights into the candidate selection that underlies the convergent evolution of the shared characteristics, especially endothermy, in birds and mammals.

## Results and discussion

### Evidence for the nocturnality of diapsid lineages

Following the MPE approach for studying the diel activity pattern of ancestral taxa ^13,20-22^, we reconstructed the diel activity patterns of our focal taxa by analyzing the adaptive evolution of 33 phototransduction genes (Supplementary Table S1) involved in the rod and cone phototransduction pathway ^23,24^ in the context of sauropsid phylogeny (Fig. 1). Positively selected genes were identified based on the branch-site model (Table 1), and the positive selection remained robust when phylogenetic uncertainties ^25-27^ (Supplementary Tables S2-3) and initial value variations of kappa and ω were considered.

**Table 1.** Positively selected genes identified based on the PAML branch-site model. For convenience, only the ω values of foreground branches are shown. Only the positively selected sites with a high posterior probability support (> 0.900) are shown.

| Taxa /Genes | Parameter estimates | 2ΔL | df | P-value | Positively selected sites |
| --- | --- | --- | --- | --- | --- |
| **Ancestral bird** |  |  |  |  |  |
| *GRK1* | $p_0=0.906$ $p_1=0.053$ $p_{2a}=0.038$ $p_{2b}=0.002$<br>$\omega_0=0.039$ $\omega_1=1.000$ $\omega_{2a}=13.983$ $\omega_{2b}=13.983$ | 9.22 | 1 | 0.002 | 20 M, 153 P, 231 C |
| *PDE6C* | $p_0=0.850$ $p_1=0.145$ $p_{2a}=0.002$ $p_{2b}=0.000$<br>$\omega_0=0.047$ $\omega_1=1.000$ $\omega_{2a}=26.734$ $\omega_{2b}=26.734$ | 6.02 | 1 | 0.014 | 49 S |
| *SLC24A1* | $p_0=0.735$ $p_1=0.250$ $p_{2a}=0.010$ $p_{2b}=0.003$<br>$\omega_0=0.050$ $\omega_1=1.000$ $\omega_{2a}=25.293$ $\omega_{2b}=25.293$ | 3.72 | 1 | 0.053 | |
| **Ancestral archosaur** |  |  |  |  |  |
| *CNGA1* | $p_0=0.871$ $p_1=0.094$ $p_{2a}=0.030$ $p_{2b}=0.003$<br>$\omega_0=0.039$ $\omega_1=1.000$ $\omega_{2a}=45.928$ $\omega_{2b}=45.928$ | 4.86 | 1 | 0.027 | 58 S, 94 A, 144 A<br>211 V, 451 R |
| *GNAT1* | $p_0=0.970$ $p_1=0.017$ $p_{2a}=0.012$ $p_{2b}=0.000$<br>$\omega_0=0.009$ $\omega_1=1.000$ $\omega_{2a}=102.241$ $\omega_{2b}=102.241$ | 5.45 | 1 | 0.019 | 147 N |
| *GUCY2D* | $p_0=0.904$ $p_1=0.091$ $p_{2a}=0.004$ $p_{2b}=0.000$<br>$\omega_0=0.046$ $\omega_1=1.000$ $\omega_{2a}=999.000$ $\omega_{2b}=999.000$ | 5.41 | 1 | 0.019 | |
| *PDE6B* | $p_0=0.896$ $p_1=0.063$ $p_{2a}=0.036$ $p_{2b}=0.002$<br>$\omega_0=0.033$ $\omega_1=1.000$ $\omega_{2a}=20.704$ $\omega_{2b}=20.704$ | 6.95 | 1 | 0.008 | 94 A, 185 T, 735 L<br>748 R |
| *RCVRN* | $p_0=0.952$ $p_1=0.024$ $p_{2a}=0.022$ $p_{2b}=0.000$<br>$\omega_0=0.020$ $\omega_1=1.000$ $\omega_{2a}=999.000$ $\omega_{2b}=999.000$ | 12.13 | 1 | 0.000 | 22 R |
| *SLC24A1* | $p_0=0.699$ $p_1=0.235$ $p_{2a}=0.049$ $p_{2b}=0.016$<br>$\omega_0=0.044$ $\omega_1=1.000$ $\omega_{2a}=4.642$ $\omega_{2b}=4.642$ | 5.14 | 1 | 0.023 | 74 T, 104 H, 149 D<br>365 N, 420 L |

2 $\Delta$ L: twice difference of likelihood values between the modified model A and the corresponding null model with the $\omega =1$ fixed in the foreground branches; df: degrees of freedom; proportion of sites and their corresponding $\omega$ values in four site classes ( $p_0$ , $p_1$ , $p_{2a}$ and $p_{2b}$ ) of the branch-site model are shown.

**Fig. 1.**
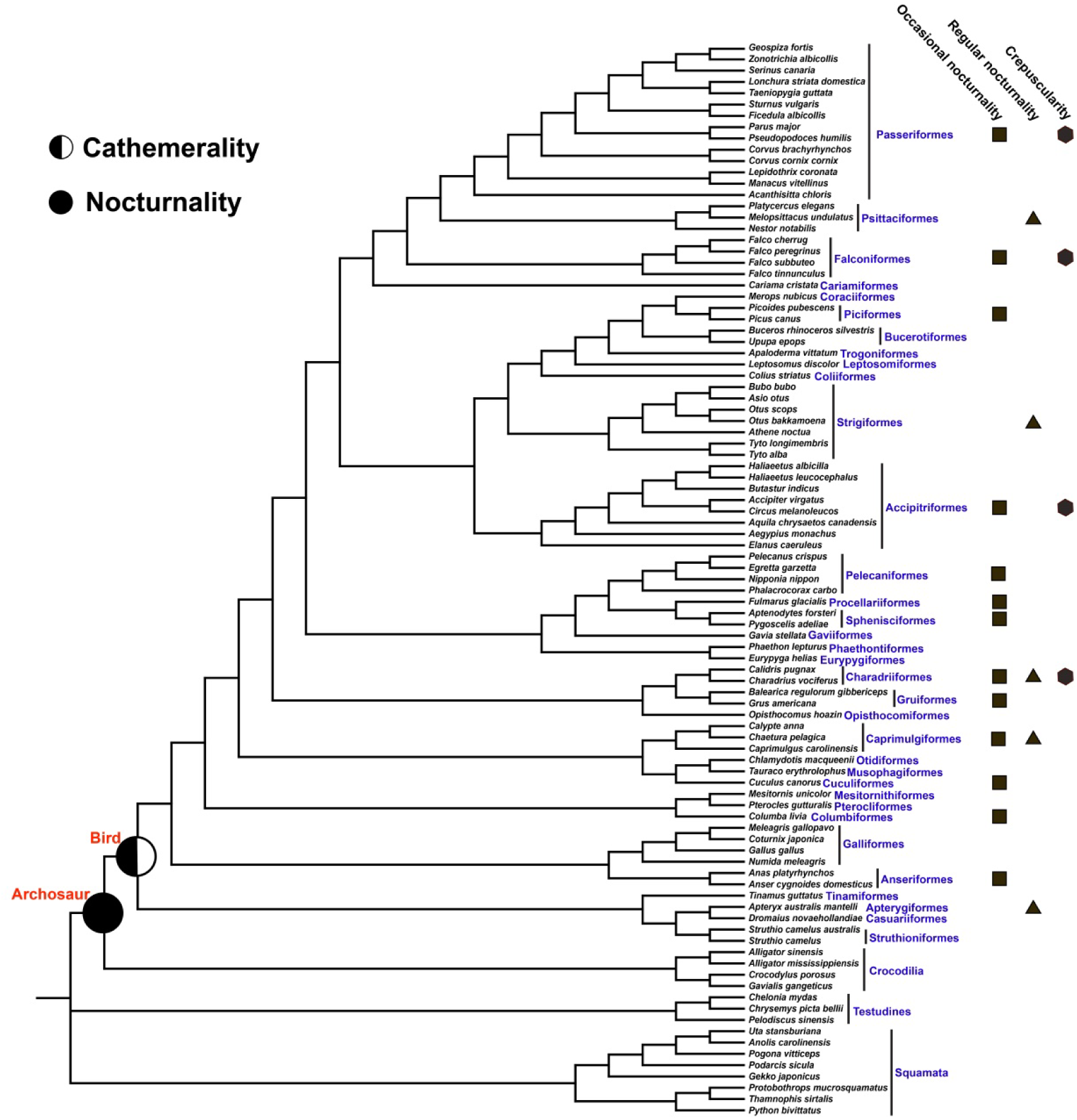
Diel activity pattern reconstruction. The phylogenetic relationships of species used in this study follows published literature ^26,68-78^ and the Tree of Life Web Project (http://tolweb.org/Passeriformes). The diel activity classification of taxonomic bird orders follows one published study ^16^. Except for five regularly nocturnal bird groups, which harbor mainly nocturnally active species, all other bird groups are considered as typically diurnal, but widely showing occasional nocturnality or engaging in partial behaviors during crepuscular periods ^16^.

We initially examined the adaptive evolution of the phototransduction genes along the common ancestor branch of living birds. One rod-expressed gene, *GRK1*, showed the most significant positive selection signal (ω = 13.983, df = 1, *P* = 0.002) (Table 1, Supplementary Fig. S1), and another rod-expressed gene, SLC24A1, showed a positive selection signal with marginal significance (ω = 25.293, df = 1, *P* = 0.053). GRK1 is rhodopsin kinase, involved in the inactivation of activated rhodopsin in rods. SLC24A1 encodes the Na^+^/Ca^2+^-K^+^ exchanger and helps to extrude free calcium in the outer segment of rods to allow the restoration of the cGMP concentration. Both genes are involved in photoresponse recovery and aid in motion detection ^28^. The finding of positive selection signals in the two rod-expressed genes suggests a particularly enhanced capability of the CALB for motion detection in dim-light conditions (e.g., nocturnality). Moreover, we found one cone-expressed gene, PDE6C, to be under positive selection (ω = 26.734, df = 1, *P* = 0.014). *PDE6C* encodes the hydrolytic subunits of cGMP phosphodiesterase (PDE6) in cones and is known to play a critical role in amplifying signals in the phototransduction cascade. The positive selection of the cone-expressed gene suggests intensified selection for increased visual acuity in bright-light environments (e.g., diurnality). Our findings of the positive selections of both the rod-expressed genes and the cone-expressed gene suggest that the CALB may have been active in both nighttime and daytime, and that they may have evolved an enhanced capability for motion detection at night.

The combined positive selection of the rod-and cone-expressed genes may also be selected for mesopic vision in crepuscular conditions, in which both rods and cones are functionally active. To test if there are shifts in their wavelengths of maximal absorption (λ_max_) towards the relatively abundant short-wavelength light in twilight, which characterizes crepuscularly active animals ^13,20^, we subsequently examined the possible spectral tuning of four cone opsin genes, *LWS, RH2, SWS2* and *SWS1*, in the CALB. Among examined critical amino acid sites (Supplementary Table S4) that are associated with spectral tuning of cone opsins ^29-32^, no critical amino acid replacement was found along the branch of the CALB, suggesting that change of the spectral tuning of the cone opsins is less likely. This may suggest that the CALB may have been mainly active in both nighttime and daytime (e.g., cathemeral) rather than being mainly active crepuscularly. This is consistent with a previous morphological study showing that the early stem bird species, *Archaeopteryx lithographica*, is somewhat intermediate in orbital characteristics between nocturnality and diurnality ^33^.

We also reconstructed the diel activity pattern of ancestral archosaur, one of the relatively distant CALB. Intriguingly, we found four rod-expressed genes, *CNGA1*, *GNAT1*, *PDE6B* and *SLC24A1*, and two genes that are rod-and cone-expressed, *GUCY2D* and *RCVRN*, which are involved in phototransduction activation or photoresponse recovery, to be under positive selection along the ancestral archosaur branch (Table 1). The positive selection of the mainly rod-expressed genes suggests enhanced visual acuity and enhanced motion detection in dim-light conditions, and hence strongly supports that ancestral archosaur was probably nocturnal, consistent with previous studies ^13,15,17^. Furthermore, these positively selected genes incorporate almost all of the principle components of the phototransduction pathway (Fig. 2), suggesting that the adaptation to nocturnality leads to a substantial modification of the visual system of the ancestral archosaur. This finding is remarkably similar to that in ancestral mammals ^13^.

**Fig. 2.**
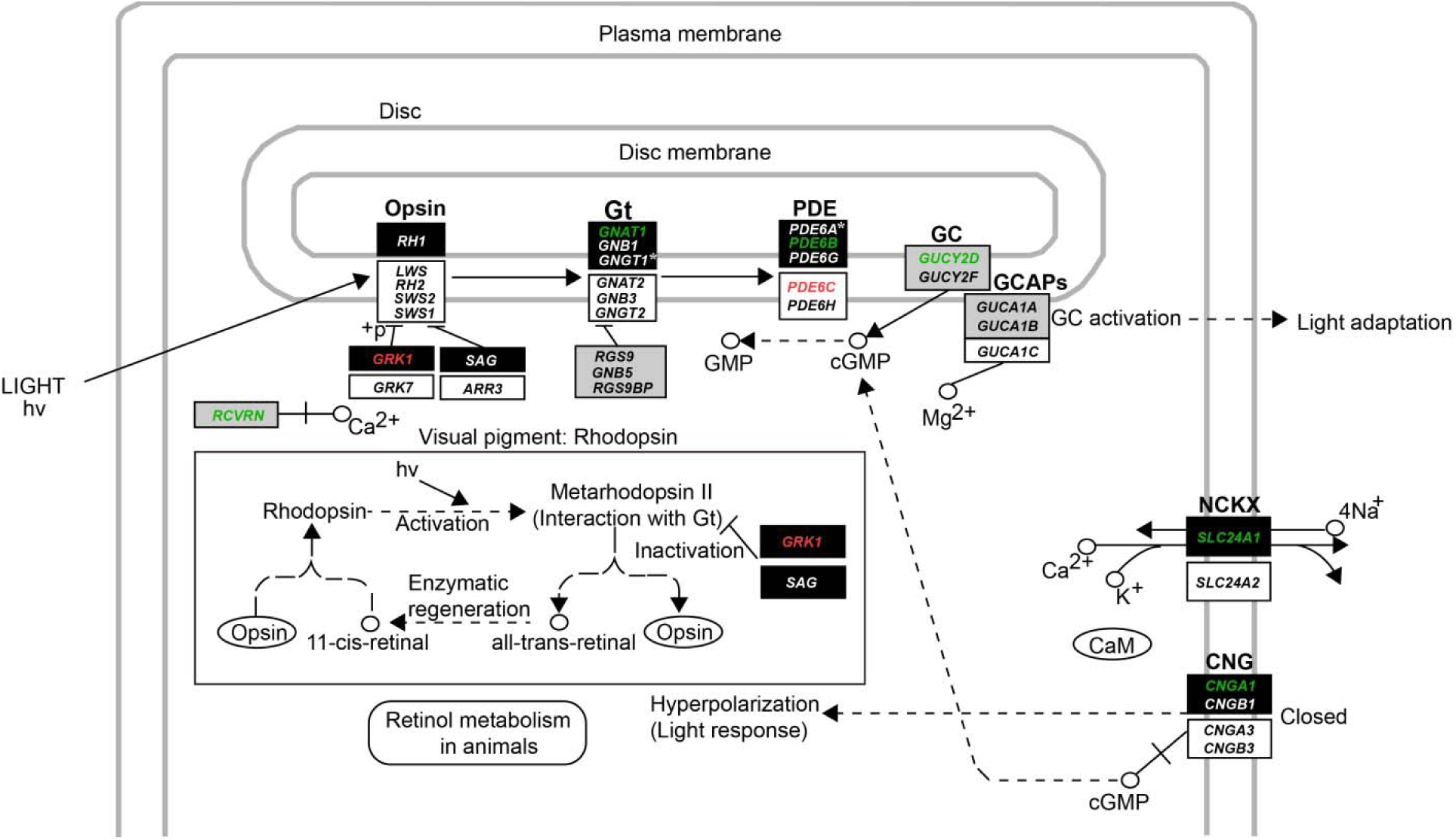
Mapping of positively selected genes on phototransduction pathway. Red and green shows the positively selected genes of the CALB and ancestral archosaurs, respectively. For convenience, both the genes involved in the rod phototransduction pathway (according to KEGG pathway: map04744) and the cone phototransduction pathway were shown. Dark rectangles, white rectangles and gray rectangles show genes that are involved in the phototransduction pathway of rods, cones and both, respectively. * Represents the losses of two rod-expressed genes, *GNGT1* and *PDE6A*, in both reptiles and birds according to previous studies ^13,20^.

Altogether, our molecular reconstruction based on the PAML results suggests that while the CALB may have been active both night and day, the ancestral archosaur was more likely nocturnal (Fig. 1). When the branch site-unrestricted statistical test for episodic diversification (BUSTED) was used to confirm the selection signals of these positively selected genes identified by PAML, one rod-expressed gene (*SLC24A1*) and one cone-expressed gene (*PDE6C*) along the ancestral bird branch and three rod-expressed genes (*CNGA1*, *PDE6B* and *SLC24A1*) along the ancestral archosaur branch were found to be under positive selection (Supplementary Table S5), providing further support for our reconstructed diel activity results of our focal taxa. In light of these robust reconstructions, our molecular results and previous studies ^13,17^ consistently suggest that the diapsid lineages may have experienced strong adaptation to nocturnality, in parallel to the synapsid lineages (e.g., early mammals) ^13^. The similar evolution of nocturnality in both diapsid and synapsid lineages should be considered as a derived result through their independent and convergent evolution, given that one previous study shows that the ancestral amniote and ancestral reptile were predominately diurnal (Fig. 3) ^13^.

**Fig. 3.**
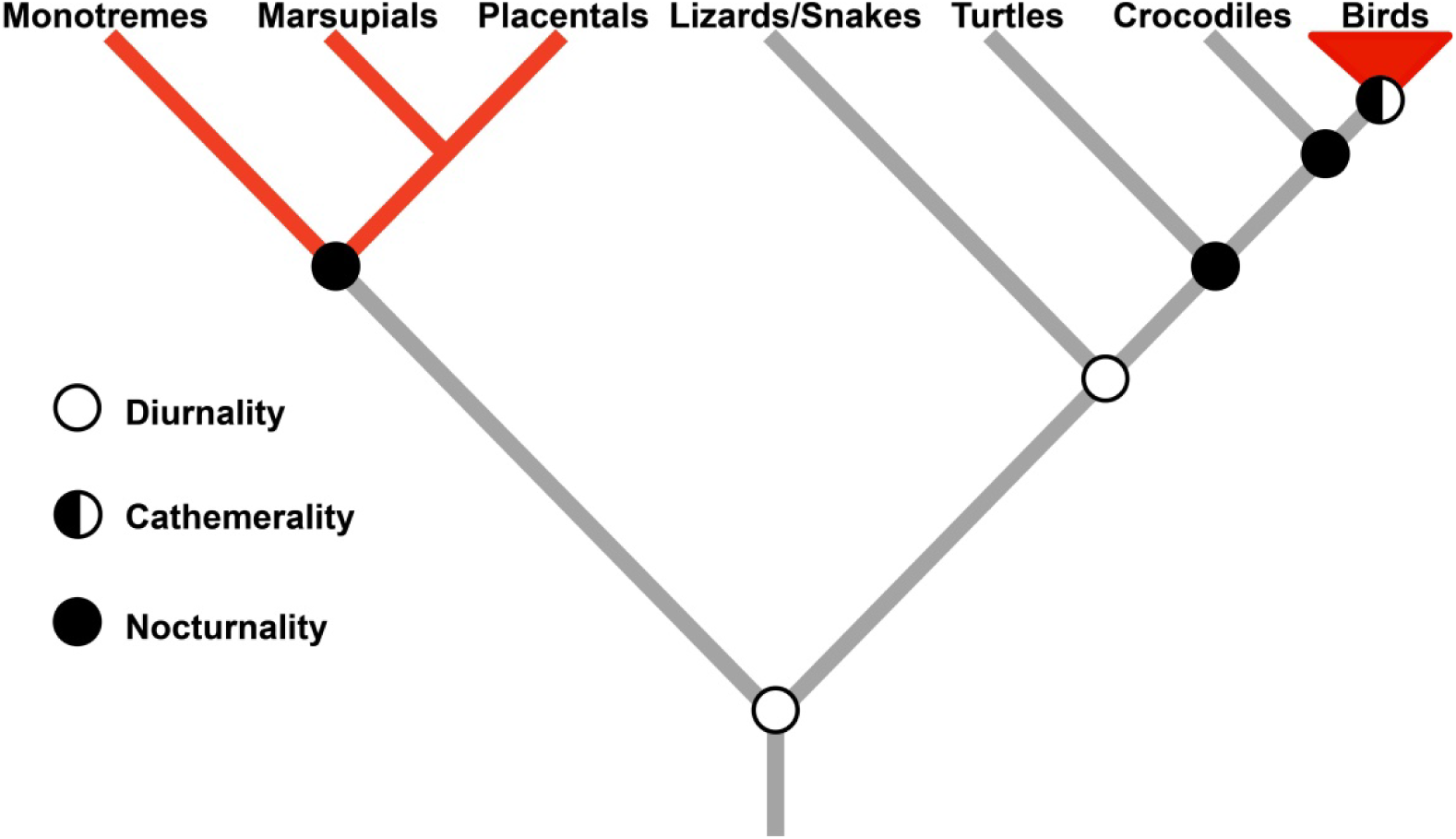
Reconstructed diel activity patterns of main groups of amniotes based on the results of this study and one previous study ^13^. Red shows endothermic groups.

### Nocturnality and endothermy evolution in diapsid and synapsid lineages

Our molecular results suggest the CALB may have experienced nocturnal periods to some extent. This nocturnality is also supported by multiple lines of morphological and behavioral evidence. 1) Theropod dinosaurs, representing one of the CALB, are reconstructed as mainly scotopic/cathemeral based on eye morphology, suggesting that the dinosaur ancestors of birds were mainly active in dim-light conditions ^17^. The discoveries of feathers (insulation) and endothermy or mesothermy in many theropod dinosaurs ^34-36^ would enable them to be active at low-temperature night, and thus are consistent with their nocturnality. 2) Though considerable controversy remains about its taxonomy, *Protoavis texensis*, which was regarded as the first bird, is characterized by owl-like large and forward-facing orbits, indicating its nocturnality ^37^. 3) Though most living birds are regarded as typically diurnal, they widely exhibit potential capability for nocturnal activities (e.g., night migration) (Fig. 1) ^16,38^, consistent with the possible nocturnal activity of the CALB. 4) Birds have the largest eyes of any of the terrestrial vertebrates ^39^, and large eyes are normally linked to increased visual acuity and are generally a character of nocturnal taxa ^16,40,41^. 5) Birds have evolved vocal communication and much more sensitive hearing than reptiles ^3^, suggesting their greater dependence on acoustic communication, which normally characterizes nocturnal species ^40,41^. Altogether, our molecular results and the behavioral and morphological evidence strongly support the nocturnality of the CALB, which challenges its pure diurnality perspectives ^15,17^.

Crompton et al. proposed that the evolution of endothermy in synapsid lineages such as mammals may have resulted from their nocturnality ^2^. Similarly, we propose that nocturnality may underlie the evolution of endothermy in diapsid lineages as well, since ancestral archosaurs, theropod dinosaurs ^17^ and ancestral birds were determined to be subjected to a similar nocturnality adaptation (Fig. 1). In support of this, previous studies demonstrate that the evolution of endothermy in diapsids may be traced back to nonavian dinosaurs ^5,34,42,43^ or even earlier to basal archosaurs ^34,44,45^. Similarly, for synapsid lineages, their nocturnality has been determined to have occurred much earlier (e.g., the Late Carboniferous almost 300 My ago) than true mammals ^46^, and their endothermy has been demonstrated to have evolved much earlier than true mammals as well, and may be traced back to the common ancestor of Neotherapsida, more than 260 My ago ^5,47-49^. In light of this, it therefore appears that the timing of the evolution of endothermy in both the diapsid and synapsid lineages coincides with the appearance of their nocturnality, supporting the nocturnality hypothesis as an explanation for their evolution of endothermy. In addition to endothermy, birds and mammals share many other characteristics, and the time for their independent evolution in the two lineages remains largely unknown. Future palaeobiological findings may help to determine the timing of their origins and help to understand the possible effects of nocturnality on their evolution.

### Nocturnality and the integrated evolution of BMSC

The diapsid (e.g., birds) and synapsid (e.g., mammals) lineages share many functionally similar characteristics, such as enhanced hearing, vocal communication, endothermy, insulation, shivering, respiratory turbinates, high basal metabolism, grinding, sustained activity, four chambered heart, high blood pressure and intensive parental care ^1-8^. The possible integrated and correlated evolution of these BMSC has long been recognized ^1,4,7,14^. Here, we attempt to explain their possible integrated evolution in the context of the adaptation to nocturnality (Fig. 4).

**Fig. 4.**
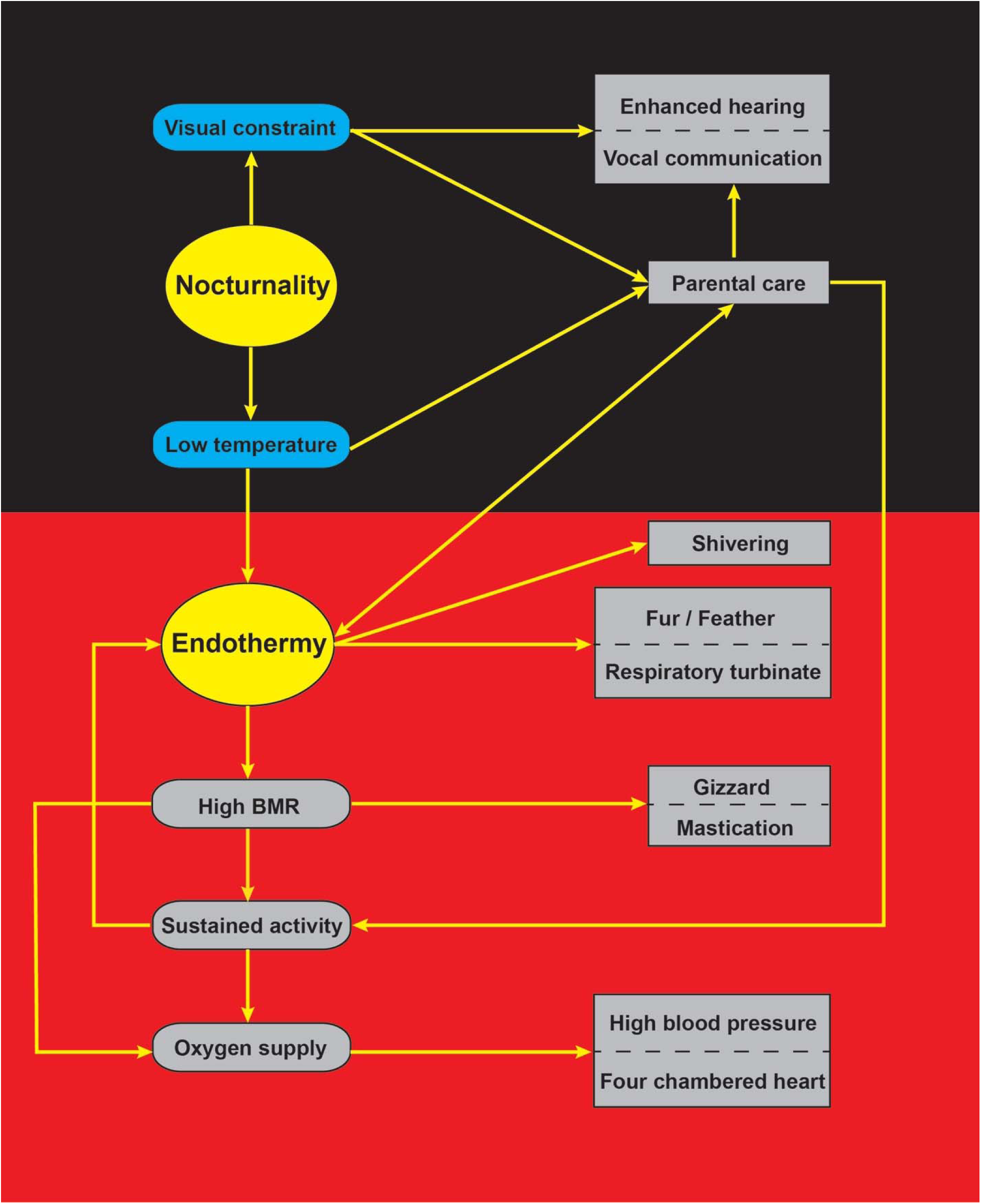
Integrated evolution of bird-mammal shared characteristics for adapting to nocturnality. Please see text for details. BMR, basal metabolic rate.

During the shift towards nocturnality of a diurnal amniote (reptile) ancestor (Fig. 3), two basal pressures: visual function constraint and low ambient temperature, may have exerted strong selection pressure for integrated evolution of the BMSC. The visual functional constraints may lead to the evolution of non-visual senses, such as enhanced hearing and vocal communication, as a result of compensatory selection pressure under dim-light conditions. In support of this, nocturnal animals generally have highly developed non-visual senses ^40,41,50^. For the low temperature selection pressure, previous studies suggest that it may lead to the evolution of endothermy, insulation (fur and feathers), respiratory turbinates or even shivering. Endothermy has long been considered as an adaptive strategy to cold temperature ^2,7,51-54^ given the temperature-dependence of many life cycle activities ^14^. Insulation (fur and feathers) and respiratory turbinates are believed to reduce heat loss ^4,5,7^, and shivering may then evolve to generate additional heat for rapid elevation of core temperature. Though its underlying thermogenesis mechanism is not fully understood, endothermy is considered to be partly linked to high basal metabolic rate (BMR) ^1^, which may then require not only accelerated assimilation through, for instance, grinding by mastication in mammals and in the gizzard in birds but also sufficient food supply ^7^. The increased food requirement may subsequently exert selection for enhanced sustained activity for efficient food acquisition. Both high BMR and sustained activity depend on an adequate oxygen supply, which may subsequently have led to the evolution of the four chambered heart, high blood pressure and other related anatomical structures ^4^.

The two selections, namely the visual function constraint and low ambient temperature associated with nocturnality, may have favored the evolution of intensive parental care as well. A juvenile of small body size has apparent disadvantages during cold nights due to its higher surface area to volume ratio, and intensive parental care would necessarily help to reduce offspring mortality ^7^. With the benefits of parental care, endothermy, hearing and vocal communication would be further strengthened to promote the offspring’s survival rate ^4^ at night. Despite this, parental care is believed to be accompanied by high costs to provision young, and this may exert additional selection pressure on sustained activity to obtain extra food ^4,8^. In addition to supporting foraging, sustained activity inevitably generates heat continuously (activity thermogenesis) ^1,55^ and may work as an important source of thermogenesis to contribute to endothermy. Activity thermogenesis, along with thermogenesis from BMR and shivering, may have worked as three principle thermogenesis sources contributing to endothermy, thereby guaranteeing the implementation of various life cycle activities under low-temperature conditions. Taken together, the evolution of BMSC is consistent with the adaptation to nocturnality (Fig. 4), and the evolution of the BMSC may be a consequence of the convergent adaption of birds and mammals to specific selections associated with nocturnality.

### Low temperature and endothermy evolution

Our results suggest that the low temperature selection pressure associated with nocturnality may have contributed to the evolution of endothermy in both diapsid and synapsid lineages. Intriguingly, although there are some controversies, low temperature is also believed to underlie the evolution of regional endothermy found in fishes (e.g., lamnid sharks, billfishes and tunas). These warm water-adapted fishes are found to frequently dive into cool water, and the origin of their endothermy was proposed to linked to multiple oceanic cooling events since the Eocene ^51-53^. Likewise, the endogenous thermogenesis of the endothermic leatherback turtle (*Dermochelys coriacea*) is found to be low-temperature dependent ^56^. Given the association between low temperature and endothermy evolution in these groups, it is possible that low temperature acts as a common factor underlying the endothermy evolution that happened independently in different vertebrate groups.

Relatively low ambient temperature seems to be an implicit condition that underlies the evolution of endotherm in different species. It is possible that low temperature also plays a role in the evolution of facultative endothermy, which was discovered in at least two reptiles during the reproductive period, the Indian python and the tegu lizard ^57,58^. The Indian python only starts to generate heat for incubation when the ambient temperature is below a critical low temperature (33°C) ^57^, which likely suggests a possible association between the appearance of facultative endothermy and the occurrence of relatively low ambient temperature. In addition, even for endothermic insects and plants, some explanations of the appearance of their endothermy have incorporated the effects of low ambient temperature ^59-61^. It is conceivable that all of these endotherms would not necessarily have evolved to generate heat themselves if their external environment constantly were similar to their elevated body temperature.

Given the possibility of the evolution of endothermy due to low temperatures (Fig. 4), parental care and sustained activity, which are popularly regarded as the initial selection pressures for the evolution of endothermy ^1,4,8^, may be only partial consequences of selection for various life cycle activities under low-temperature conditions, instead of the sole or even the initial selective target. Regarding the two popular views, criticisms can be found elsewhere ^4,8,62^, while one additional issue is that no explanation is provided to interpret why only avian and mammalian lineages were selected for sustained activity or intensive parental care but other living reptiles have not done likewise. Moreover, even for the biological functions of endothermy, it would be biased to only emphasize one of its many potential biological functions, leading to many theoretical models applicable only to specific groups ^9,11^. For instance, neither the aerobic capacity model ^1^ nor the parental care model ^4^ explain the evolution of regional endothermy found in fishes, which are known to have evolved for other possible functions rather than for sustained activity or parental care ^51,53^. In fact, for all different endothermic organisms found to date, endothermy is involved in diverse functions, spanning all possible life cycle activities related to locomotion, sensory communication, foraging, digestion, reproduction, breeding, antipredation and parental care ^1,4,14,51,53,59^. A conceptually unifying model accounting for all of these diverse functions of endothermy in the context of low-temperature environments would be more reasonable to explain the appearance of endothermy in diverse organisms. In light of this, we propose that endothermy may evolve as an adaptive strategy to enable organisms to efficiently implement various life cycle activities (e.g., parental care and sustained activity) under relatively low-temperature environments.

### Endothermy evolution and the habitat shift from high-temperature environments to low-temperature environments

The diel activity changes of both diapsid and synapsid lineages indicate that they may have experienced a habitat shift from high-temperature environments (day) to relatively low-temperature environments (night) (Fig. 3), which may underlie the evolution of their endothermy. Similarly, it is suggested that the evolution of endothermy in diverse fishes may be linked to their habitat change from warm-water habitat to cold-water habitat ^51-53^. Likewise, for leatherback turtles, the appearance of their endothermy is associated with their habitat change from tropical nesting sites to cold northern foraging grounds ^56^. For the facultative endothermic reptile, Indian python (*Python molurus*) ^57^, one molecular phylogeographic study reveals a possible origin of the genus *Python* in West Africa followed by dispersal to Europe and Asia ^63^, suggesting that the evolutionary origin of the Indian python may have experienced a habitat shift from high temperature areas (tropical climate) to relatively low temperature areas (subtropical/temperate climate). The other facultative endothermic reptile, the tegu lizard (*Salvator merianae*) ^58^, might have experienced a similar high-to-low temperature habitat shift, as it extends geographic distribution from tropical, subtropical to relatively low-temperature temperate climates compared with its closest relatives (*Salvator duseni, Salvator rufescens*) ^64^.

In light of these findings, it appears that all known endothermic vertebrates may have been subjected to a habitat shift from high-temperature to relatively low-temperature environments during their evolutionary histories. This specific habitat shift may be an important means to stimulate the evolution of endothermy in vertebrates. Endothermy would be selectively favored in low-temperature environments for optimum activity of enzymes and other molecules that had been adapted to previous relatively high-temperature conditions. If this is true, we may expect that endothermy would be more likely to be found in taxa with their biogeographical histories involving a habitat shift from high-temperature environments to relatively low-temperature environments. This pattern remains to be tested in relevant endothermic vertebrates, insects and plants ^59,61^.

Endothermy may not be the sole strategy to adapt to low-temperature environments. Apparently, some ectothermic animals, for instance, geckos and cold-water fish, are capable of being active in cold-temperature environments, suggesting the existence of alternative physiological or behavioral strategies to respond to cold temperatures. In addition to low temperature, many other possible factors such as biogeographic history, food availability, body size, and selection intensity may also affect the evolution of endothermy. Future study through a comparative and integrated analyses of possible factors between endotherms and their phylogenetically closest ectotherms should help to explain why endothermy is only restricted to some specific groups.

## Conclusion

This study provides strong evidence for the nocturnality of the diapsid lineages, paralleling that of the synapsid lineages. Given their nocturnality, an integrated perspective on the evolution of BMSC as a convergent adaptation to nocturnality is proposed. Moreover, after summarizing our findings and relevant empirical studies on the evolution of endothermy, low temperature is suggested as a possible common factor underlying endothermy evolution in vertebrates. Given the significance of low temperature in endothermy evolution, a conceptually unifying ecological model of endothermy evolution with an emphasis on low temperature is proposed. We reason that endothermy may evolve as an adaptive strategy to enable organisms to effectively implement various life cycle activities under relatively low-temperature environments, which happens during a habitat shift from a high-temperature environment to a relatively low-temperature environment.

## Materials and Methods

### Taxa covered

The retinal transcriptome sequencing data of 17 bird species of which 15 were published previously by our lab ^20^ and two newly generated in this study, along with all of available genome data of birds and reptiles in GenBank were used in this study (Supplementary Table S1). In total, 95 species were used, of which 80 bird species from 34 orders, representing the majority of living bird orders (34/39) ^65^, were covered. During our initial sequence analyses, we found several of our focal phototransduction genes missing from paleognath genome data, and thus two species, the ostrich (*Struthio camelus*) and the emu (*Dromaius novaehollandiae*), were selected to be subjected to retinal transcriptome sequencing.

### Sampling, RNA isolation and cDNA library construction

One ostrich (three months old) and one emu (six months old) were obtained with permission from an artificial breeding company (Quanxin, Daqing). Active individuals were transported to the laboratory on December 2, 2017 and maintained under natural light:dark conditions with food and water provided. The ostrich and emu were sacrificed after 24 hours and their retinal tissues were dissected during the daytime (3 pm to 4 pm) and were preserved in RNA-locker (Sangon Biotech, Shanghai), flash frozen in liquid nitrogen, and then stored at −80°C until processed for RNA isolation one week later. The experimental procedures were performed in accordance with an animal ethics approval granted by Northeast Normal University. All experimental procedures carried out in this study were approved by the National Animal Research Authority of Northeast Normal University, China (approval number: NENU-20080416) and the Forestry Bureau of Jilin Province of China (approval number: [2006]178).

Total RNA isolation was performed using TRIzol Reagent (Invitrogen Life Technologies), following the manufacturer’s protocol and instructions. RNA degradation and contamination was monitored on 1% agarose gels. RNA purity was checked using the NanoPhotometer® spectrophotometer (IMPLEN, CA, USA). RNA concentration was measured using the Qubit® RNA Assay Kit in a Qubit®2.0 Flurometer (Life Technologies, CA, USA). RNA integrity was assessed using the RNA Nano 6000 Assay Kit of the Agilent Bioanalyzer 2100 system (Agilent Technologies, CA, USA). For cDNA library construction, a total amount of 3 μg RNA per sample was used as input material for the RNA sample preparations. The cDNA library was generated using NEBNext®Ultra™ RNA Library Prep Kit for Illumina® (NEB, USA) according to the manufacturer’s protocol. Specifically, mRNA was purified from total RNA using poly-T oligo-attached magnetic beads. The enriched mRNA was fragmented into small pieces and was then used for the syntheses of the first and the second strands of cDNA. The cDNA was purified with the AMPure XP system (Beckman Coulter, Beverly, MA, USA), and then 3 μl USER Enzyme (NEB, USA) was used with size-selected, adaptor-ligated cDNA at 37°C for 15 min followed by 5 min at 95°C before PCR. Then, PCR was performed, PCR products were purified (AMPure XP system) and library quality was assessed on the Agilent Bioanalyzer 2100 system. Paired-ending sequencing of the cDNA library was performed using Illumina HiSeq X Ten (Biomarker Technology Co., Beijing).

### Data filtering and de novo assembly

In total, 10.48G bases and 12.05G bases were generated for ostrich and emu, respectively. Raw data were filtered by removing reads containing adapter, reads containing ploy-N and low quality reads, and clean reads with high quality were obtained and used for de novo assembly. De novo assembly was performed using Trinity ^66^ with default parameters except that min_kmer_cov =2 was used, ensuring that only k-mers with at least 2x coverage were used. The longest transcript was used to yield unigenes, and the unigenes longer than 200⟊bp were retained for further analyses. The transcriptome sequencing data of the two species have been deposited into the National Center for Biotechnology Information Sequence Read Archive database under accession number (SRP134220).

### Sequence abstraction and alignment

The 33 phototransduction genes involved in both the rod phototransduction pathway and the cone phototransduction pathway were used in this study ^23^. To extract the coding sequences from the ostrich and the emu unigene pools, the coding sequences of *Gallus gallus* downloaded from GenBank were used as query sequences to blast against the unigene pools of the two species by using Blastn. The unigene sequences obtained were then annotated by blasting against the NCBI nr/nt database using the online Blastn, and only the unigene sequences with the same gene annotation as that of the query sequences were used for subsequent analyses. In addition to these two species, the coding sequences of the 33 genes from our previously published 15 bird species were also included ^20^. Additionally, we downloaded the longest transcript sequences of our target genes from all other birds and reptiles with genome data available in GenBank. Gene sequences were aligned using webPRANK (http://www.ebi.ac.uk/goldman-srv/webprank/), and individual sequences with low identities, long indels, multiple ambiguous bases Ns, and/or too short a length were removed or replaced by other relevant transcript variants. After this pruning, high-quality alignments were constructed, and their translated protein sequences were blasted against the non-redundant protein sequence database to confirm the correctness of sequence cutting.

### Positive selection analyses

We used the branch model and branch-site model implemented in the Codeml program of PAML ^67^ for positive selection analyses. Upon analysis, we initially constructed an unrooted species tree following published studies ^26,68-78^ and the Tree of Life Web Project (http://tolweb.org/Passeriformes). The phylogenetic relationships among bird orders followed one genome-level study ^26^. We used a codon-based maximum likelihood method to estimate the ratio of non-synonymous to synonymous substitutions per site (dN/dS or ω), and employed likelihood ratio tests (LRT) to determine statistical significance. A statistically significant value of ω ⟊ > ⟊1 suggests positive selection.

#### Branch model

We used a two-rate branch model to detect possible positive selection signals along the ancestral bird branch and the ancestral archosaur branch. For analyses, the two branches were respectively labelled foreground branches and others were treated as background branches. In the two-rate branch model, ω is assumed to be different between foreground branches and background branches, and its goodness of fit was analyzed using the LRT, in which the two-rate branch model was compared with the one-rate branch model that assumes a single ω value across the tree. If a statistically significant value of ω ⟊ > ⟊1 in a foreground branch was detected, the two-ratio branch model was then compared with the two-ratio branch model with a constraint of ω = 1 to further determine whether the ω ⟊ > ⟊1 of the foreground branch was statistically significant.

#### Branch-site model

We used a branch-site model (Test 2) to detect positively selected sites for our focal branches (foreground branches). In this model, four classes of sites are assumed. Site class 0 (0 < ω_0_ < 1) and site class 1 (ω_1_ = 1) represent evolutionarily conserved and neutral codons, respectively, for both background branches and foreground branches. Site classes 2a and 2b represent evolutionarily conserved (0 < ω_0_ < 1) and neutral (ω_1_ = 1) codons, respectively, for background branches but are allowed to be under positive selection (ω_2_ > 1) for the foreground branches. Test 2 compares a modified model A with its corresponding null model with ω = 1 constrained to determine the statistical significance. The empirical Bayes method was used to detect positively selected sites.

### Robustness test of positive selection genes

The positively selected genes detected were further subjected to a robustness test. First, to detect the dependency of our positive selection results on the initial value variations of kappa and omega, we used two different initial values of kappa (kappa = 0.5, 3.0) and of omega (ω = 0.5, 2.0) for our positive selection analyses. In total, four independent runs were conducted for each of the positively selected genes found. Moreover, we examined the effects of phylogenetic uncertainties on our positive selection results. So far, considerable phylogenetic inconsistencies among different bird orders have been found. To examine the phylogenetic uncertainty effects, we used three different phylogenetic relationships of bird orders ^25-27^ for our positive selection analyses. For convenience, the three different phylogenies were referred to as Hackett’s tree, Jarvis’s tree and Prum’s tree in our study.

For our results robustness, we also employed BUSTED, implemented in the HyPhy software (version 2.2.4) ^79,80^, to confirm the positive selection signals of those positively selected genes identified by PAML. Unlike PAML, in which only positive selection in foreground branches is allowed, BUSTED allows the occurrence of positive selection in both foreground and background branches (unconstrained branch-site model). The unconstrained model fit is compared with the branch-site model that disallows ω > 1 among foreground branches (null model), and then LRT was used to determine statistical significance.

### Spectral tuning analyses

We analyzed possible spectral tuning (i.e., the shifts of wavelengths of maximal absorption) of four cone-expressed opsins, *LWS*, *RH2*, *SWS2* and *SWS1*, along the ancestral bird branch and the ancestral archosaur branch. For this, we reconstructed ancestral amino acid sequences using amino acid-based marginal reconstruction implemented by the empirical Bayes approach in PAML 4.8a to determine the amino acid replacements along the branches, considering that the spectral tuning of these cone opsins is controlled by specific critical amino acid replacements ^29-32^. To test for result consistency, two different amino acid substitution models, JTT and Poisson, were used. The two models have different assumptions about amino acid substitution rates. The model JTT assumes different substitution rates of different amino acids, while the Poisson model assumes the same substitution rate of all amino acids. With the reconstruction of ancestral sequences for internal nodes, the critical amino acid replacements that are associated with the spectral tuning of cone opsins along branches can be inferred. To determine the numbering of the critical amino acid replacements in our sequences, we employed webPRANK to align our sequences against the bovine rhodopsin sequence (NP_001014890). The effects of the critical amino acid replacements on the shifts of the wavelengths of maximal absorption (Δ λ) of the four cone opsins were analyzed following published studies ^29-32^.

## Acknowledgements

We thank Lin Chen, Yuanqin Zhao and Li Gu for helping tissue sampling. This research was supported by the National Natural Science Foundation of China (grant number, 31770401) and the Fundamental Research Funds for the Central Universities.

## Author Contributions

Y. W. designed research, performed analyses and wrote the paper. H. W. revised the manuscript. All authors gave final approval for publication.

## Competing interests

We have no competing interests.

## Data and materials availability

All data needed to evaluate the conclusions in the paper are present in the paper and/or the Supplementary Materials. Additional data available from authors upon request.

## Supplementary Materials

Table S1-S5

Fig S1

